# Associated Bacteria, Particularly *Mucilaginibacter* Species, Inhibit Sheath Formation in *Nostoc commune*

**DOI:** 10.64898/2026.10.14.759064

**Authors:** Masaki Ihara, Sora Misaizu, Kaede Tomiyama, Maeka Shiroyama, Yusuke Sakai, Takamichi Nakamura, Hiroyuki Kakenaka

## Abstract

The terrestrial cyanobacterium *Nostoc commune* fixes atmospheric nitrogen and exhibits remarkable desiccation tolerance, making it a promising organism for biomass production on land unsuitable for conventional agriculture. However, reliable artificial cultivation of *N. commune* remains difficult because hormogonium formation cannot yet be induced reproducibly and mechanically prepared trichome fragments often fail to develop into sheath-enclosed colonies. In this study, we examined factors affecting sheath formation in short non-motile trichome fragments, referred to here as nas-trichomes. Microscopic observation revealed two distinct developmental outcomes: a curling type, in which trichomes formed sheaths and developed into compact colonies, and an elongation type, in which trichomes extended without forming a distinct sheath. Curling-rich cultures were obtained only from colonies subjected to drying and rehydration, and early bacterial proliferation around trichomes was negatively associated with subsequent sheath formation. Microbial community analysis showed that *Mucilaginibacter* was enriched in elongation-rich cultures. Co-culture experiments further demonstrated that *M. segetis, M. oryzae, M. jinjuensis*, and *M. xinganensis* significantly inhibited sheath formation at high bacterial density. These results indicate that associated bacteria, particularly specific *Mucilaginibacter* species, can inhibit sheath formation in *N. commune* and suggest that controlling early bacterial proliferation is essential for establishing reliable cultivation methods.

## Introduction

*Nostoc commune* is a filamentous terrestrial cyanobacterium that forms large macroscopic colonies enclosed within a common polysaccharide-rich sheath. Numerous trichomes are embedded within this sheath and aggregate to form gelatinous, ball-like colonies that are visible to the naked eye (Briones-Nagata et al. 2007). Owing to this distinctive colonial architecture and its remarkable environmental resilience (Inoue-Sakamoto et al. 2018, Sand-Jensen and Jespersen 2012, Fukuda et al. 2008, Wright et al. 2005, Tamaru et al. 2005, Satoh et al. 2002, Bazzichelli et al. 1985), *N. commune* has attracted increasing attention as a biological resource that may contribute to sustainable biomass production.

As an oxygenic photosynthetic cyanobacterium capable of fixing atmospheric CO_2_, and as a diazotroph capable of biological nitrogen fixation through heterocysts, *N. commune* can convert atmospheric carbon and nitrogen into biomass with minimal dependence on fossil-fuel-derived inputs. These traits make it a promising candidate for the production of low-carbon fertilizers and animal feed (Katoh et al. 2012). In addition, its biomass contains bioactive constituents, including extracellular polysaccharides and proteins, suggesting potential applications as a functional food resource (Watanabe et al. 2007, Ninomiya et al. 2011, Itoh et al. 2013, Sasaki et al. 2013, Wang 2026, Li and Guo 2018).

Despite proliferating extensively across a wide range of terrestrial habitats in nature, a reliable and reproducible method for the artificial cultivation of *N. commune* has yet to be established. A major factor underlying this gap is that, although *N. commune* reproduces and disperses through the release of motile filaments called hormogonia, the process governing hormogonia release cannot currently be controlled artificially.

During routine laboratory observation of *N. commune* colonies, we have occasionally observed the sheath rupture, followed by fragmentation of the trichome at sites flanking heterocysts, and the subsequent emergence of hormogonia—short filaments composed of a dozen or so cells— swimming out through the resulting opening in the sheath. To our knowledge, however, no study has specifically reported on hormogonia release in *N. commune*.

Consequently, current cultivation approaches instead rely on mechanically homogenizing colonies to obtain short trichome fragments (hereafter, “nas-trichomes”), which are then cultured on agar medium. Although these nas-trichomes are occasionally able to develop into spherical colonies (Katoh et al. 2012), cultures frequently die before reaching this stage, and the cause of this culture failure remains unknown.

The sheath of *N. commune* has generally been regarded as an adaptation that confers tolerance to desiccation and other environmental stresses (Inoue-Sakamoto et al. 2018, Hill et al. 1994, Ehling-Schulz et al. 2002, Tamaru et al. 2005, Wright et al. 2005, Sand-Jensen and Jespersen 2012). We hypothesized that the sheath may also play an important role in defense against microbial attack. Consistent with this idea, we observed that *N. commune* cells lacking a sheath died rapidly. In addition, when intact colonies were soaked in water for an extended period, the sheath dissolved (Morsy et al. 2008), the enclosed cells died, and bacteria consistently aggregated around the dead cells. Bacterial predation is known to pose a serious threat to terrestrial cyanobacteria (Bauer and Forchhammer 2021, Bethany et al. 2022).

Based on these observations, we investigated whether associated bacteria, particularly bacteria enriched in sheath-deficient cultures, inhibit sheath formation and thereby affect the survival and colony formation of *N. commune* trichomes. We further examined whether candidate bacteria can reproduce sheath-formation inhibition in co-culture, with the aim of identifying microbial factors that limit reliable artificial cultivation.

## Materials and Methods

### Strains

*Nostoc commune* colonies used in this study were collected from the banks of the Mibu River in Ina City, Nagano Prefecture, Japan (137.6°E, 35.5°N). *Mucilaginibacter oryzae* (NBRC 106410), *M. jinjuensis* (NBRC 108856), *M. xinganensis* (NBRC 1100384), *M. pallidiroseus* (NBRC 113666), *M. achroorhodeus* (NBRC 113667), and *Ideonella sakaiensis* (NBRC 110686) were obtained from the NITE Biological Resource Center (NBRC). *Mucilaginibacter conchicola* (JCM 32787), *M. inviolabilis* (JCM 34116), and *M. segetis* (JCM 34284) were obtained from the Japan Collection of Microorganisms at RIKEN BioResource Research Center. Lyophilized bacterial cultures were rehydrated according to the suppliers’ instructions. *Escherichia coli* HB101 was used as the *E. coli* strain. *Lactiplantibacillus plantarum* strain k4-9 was kindly provided by Sachi Tanaka (Shinshu University) (Tanaka et al. 2024).

### Media

The composition of BG11_00_ medium used for culturing *N. commune* is shown in Table S1. All reagents listed in Table S1, except FeSO4·7H2O, were dissolved in deionized water (DIW) with thorough stirring and autoclaved at 121°C for 20 min. Before use, filter-sterilized FeSO4·7H2O solution was added to the autoclaved medium. BG11_00_ agar medium was prepared in the same manner, except that an autoclaved agar suspension (Nacalai Tesque, Inc., Kyoto, Japan) was added after autoclaving to a final concentration of 1.5% (w/v). For rehydration of dried *N. commune* colonies, DIW containing 1.5 ppm sodium hypochlorite (NaClO; Nacalai Tesque, Inc.) was used and is hereafter referred to as hypochlorite solution. *Escherichia coli* was cultured in LB medium prepared by dissolving 20 g of LB powder (Nacalai Tesque, Inc.) in 1 L of water and autoclaving at 120°C for 20 min. Lactic acid bacteria were cultured in MRS medium prepared by dissolving 55 g of Difco Lactobacilli MRS Broth powder (BD, Franklin Lakes, NJ, USA) in 1 L of water and autoclaving at 121°C for 20 min. *Nostoc commune* polysaccharide agar medium, used to isolate bacteria potentially affecting sheath formation, was prepared as follows. Approximately 5 g of wet *N. commune* colonies was suspended in 10 mL of DIW and homogenized with a 10 cm3 glass homogenizer equipped with a motorized stirrer (HK-1; AS ONE Corporation, Osaka, Japan) until no visible clumps remained. The homogenate was brought to 125 mL with DIW and autoclaved. Autoclaved agar solution containing 3.75 g agar powder per 125 mL was then mixed with the homogenate to prepare *N. commune* polysaccharide agar medium.

**Table S1.**
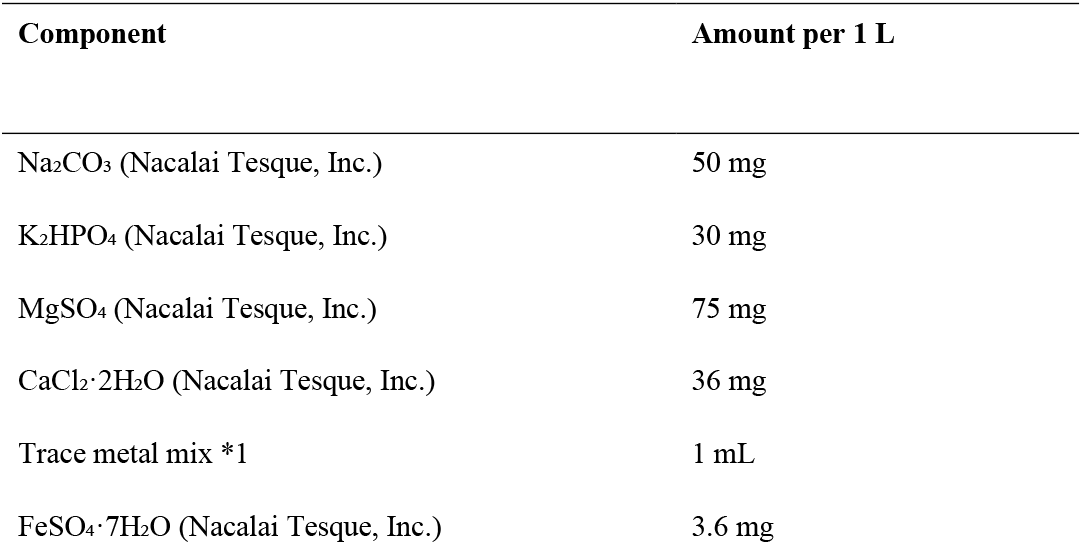
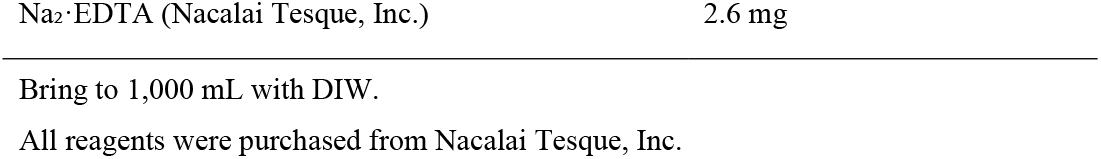
Composition of BG11_00_ medium used for cultivation of *Nostoc commune*.

*1 Trace metal mix composition, prepared in 1,000 mL of RO water

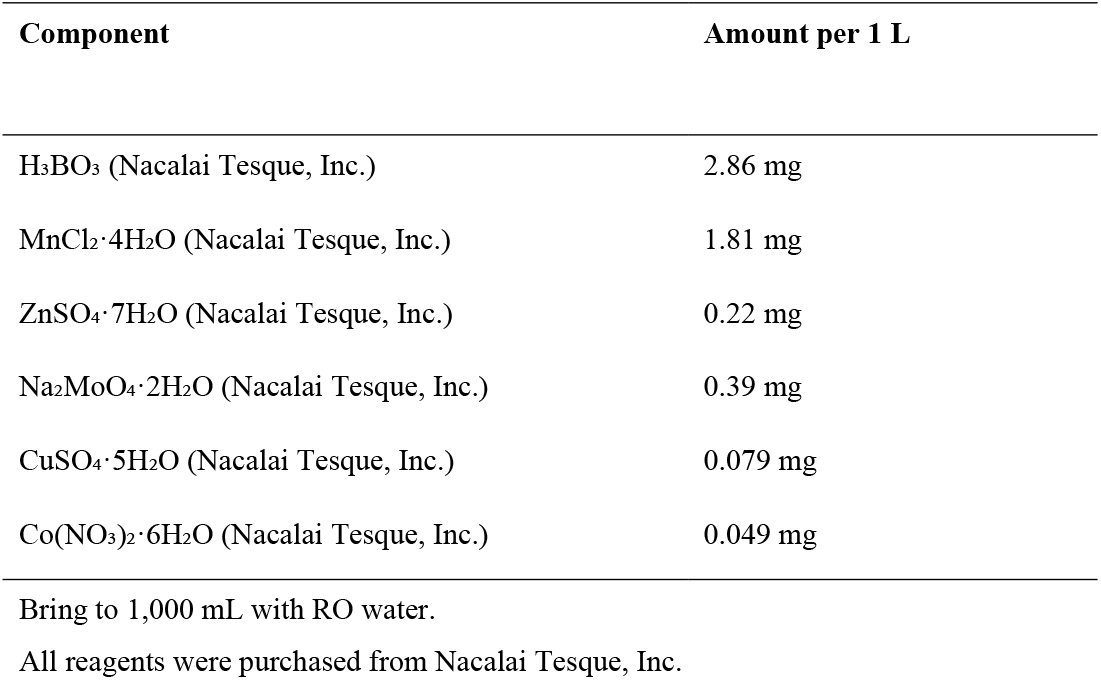

### Cultivation of *N. commune*

Collected *N. commune* colonies were immersed in DIW and rehydrated for approximately 12 h at 20–25°C. Soil, leaves, and other debris were then removed, and the colonies were rehydrated for an additional 12 h. This rehydration–washing cycle was repeated three times in total. The washed colonies were placed on a mesh with approximately 1-mm openings in an incubator set at 35°C, and air passed through a cotton filter was blown upward through the mesh for 24 h to dry the colonies. The dried colonies were immersed in 1.5 ppm hypochlorite solution and rehydrated for 12 h in the dark at 20–25°C. The rehydrated colonies were rinsed three times with DIW, cut into approximately 0.5–1-cm pieces, and transferred to a 15-mL centrifuge tube containing 10 mL of DIW until the total volume reached approximately 11 mL. The contents were transferred to a 10 cm3 glass homogenizer, and the pestle was connected to a motorized stirrer. Colonies were disrupted at 3,000 rpm with 20 reciprocating strokes at approximately 2 s per stroke. Sheath fragments were removed by passing the homogenate through a 40-µm pluriStrainer (pluriSelect Life Science, Leipzig, Germany), and the filtrate was then passed through a 10-µm pluriStrainer to recover *N. commune* trichomes. Because polysaccharides caused clogging, vacuum filtration was performed while the liquid on the 10-µm filter was gently pipetted. Sterile water was then added, and filtration was repeated four times. The material retained on the 10-µm filter was collected and centrifuged at 3,200 rpm for 5 min. The supernatant was discarded, and the trichome pellet was resuspended in 500 µL of liquid BG11_00_ medium. These artificially severed, non-motile trichomes are hereafter referred to as nas-trichomes. Ten microliters of the nas-trichome suspension was loaded onto a C-Chip hemocytometer (NanoEntek, Seoul, Korea), and nas-trichomes were counted under a biological microscope. The suspension was diluted with DIW to 10–30 nas-trichomes per 10 µL, and 10 µL of the diluted suspension was added to each well of a 96- or 384-well plate containing 50 µL of BG11_00_ medium. Plates were incubated at 25°C under white LED illumination at 50 µmol photons m−2 s−1 in a temperature-controlled chamber. After 5– 10 days, wells in which >90% of trichomes grew with curling and formed a sheath-enclosed tangle (Fig. 1B) were designated curling-rich wells, whereas wells in which >90% of trichomes elongated without forming a distinct sheath (Fig. 1D) were designated elongation-rich wells.

**Fig. 1.**
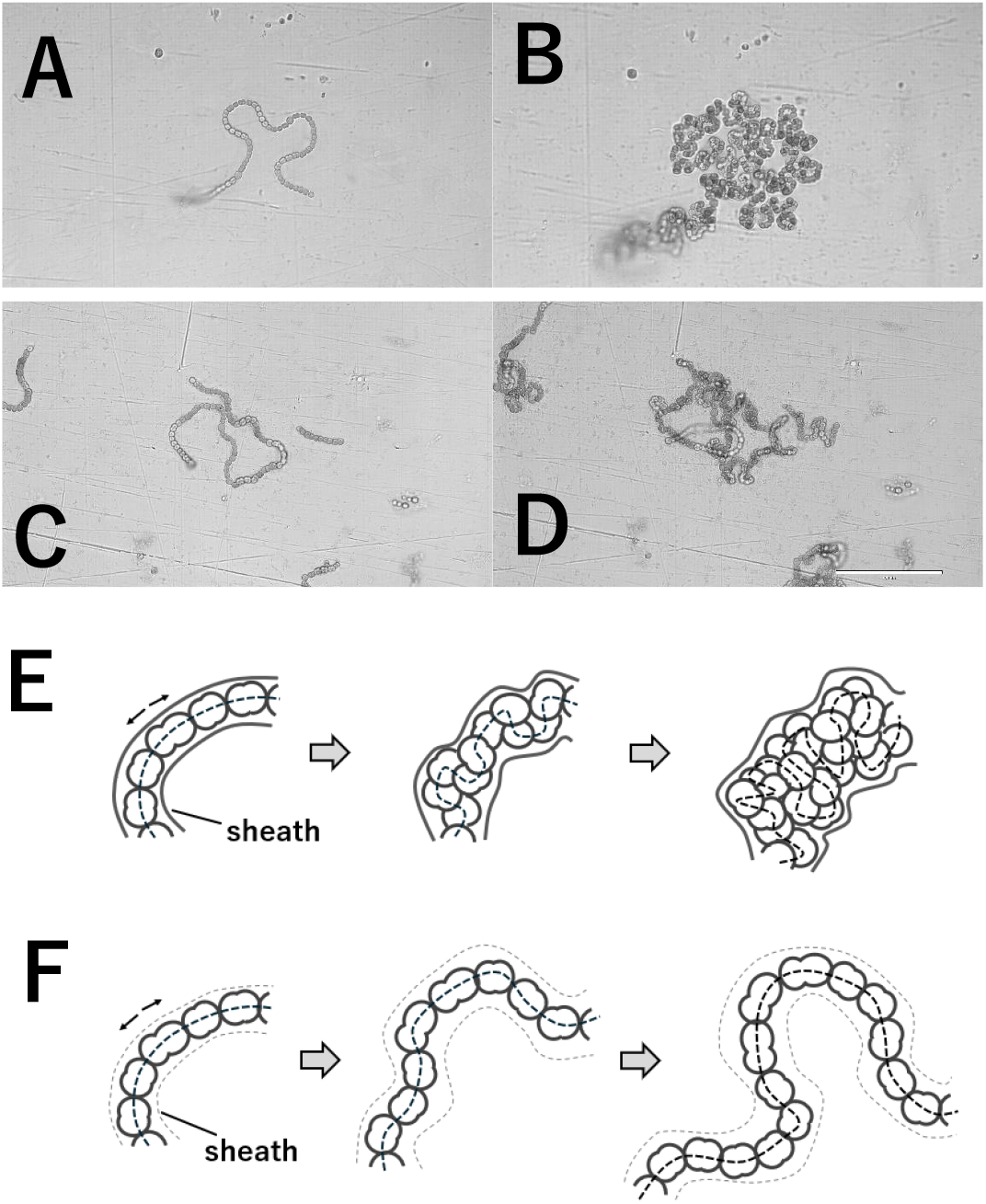
Morphological changes in *N. commune* trichomes during culture. (A) Nas-trichomes prepared from colonies subjected to drying and rehydration treatment, 3 days after the start of cultivation. (B) The same field as in (A), 5 days later, corresponding to day 8 of cultivation. Cells within the trichome increased in number through division, but spreading was restricted by the sheath, resulting in microcolony formation. (C) Nas-trichomes prepared from untreated colonies, 3 days after the start of cultivation. (D) The same field as in (C), 5 days later, corresponding to day 8 of cultivation. Trichomes without a distinct sheath elongated through cell division. Numerous particles, presumably bacteria, were observed around the trichome. (E) Schematic illustration of proliferation of a sheathed trichome. Filamentous trichomes are depicted as serially connected cells enclosed within a sheath. Dashed lines indicate the approximate central axis of connected cells, and the arrow indicates the direction of cell division and proliferation. Sheathed trichomes proliferated in a compact, ball-like manner. (F) Schematic illustration of elongation of a trichome without an obvious sheath. A faint, unclear sheath is indicated by light dashed lines, although it was not clearly visible microscopically. Bold dashed lines indicate the trajectory of cell connections, and the arrow indicates the direction of cell division and proliferation. Trichomes lacking a distinct sheath elongated in a filamentous manner. Scale bar = 100 µm.

### 16S rRNA Analysis of *N. commune* Culture Fluid

Nas-trichome suspensions were prepared individually from 50 natural colonies collected from the Mibu River, and each suspension was dispensed into 96 wells of a 384-well plate. Microscopic observation confirmed that each well contained 30–80 trichomes. A set of wells containing nas-trichomes derived from the same colony is referred to as a group. After 10 days of cultivation, the numbers of curling-rich and elongation-rich wells were counted for each group. Groups containing more than 40 curling-rich wells were designated S(+) groups, whereas groups containing more than 40 elongation-rich wells were designated S(−) groups. Nas-trichomes were recovered from curling-rich wells in S(+) groups and from elongation-rich wells in S(−) groups, pooled within each group, and stored at −25°C until DNA extraction. DNA was extracted using the Extrap Soil DNA Kit Plus Ver. 2 (BioDynamics Laboratory Inc., Tokyo, Japan). DNA concentrations were measured using a Qubit Fluorometer with the Qubit dsDNA HS Assay Kit (Thermo Fisher Scientific, Waltham, MA, USA). Library preparation and 16S rRNA sequencing were outsourced to FASMAC Co., Ltd. (Kanagawa, Japan).

### Purification of *N. commune*

Purification of *N. commune* nas-trichomes was attempted using antibiotic treatments based on the method of Kan and Pan (2010). Nas-trichomes were cultured on BG11_00_ agar medium supplemented with ampicillin (Amp; 500 µg/mL), cefotaxime (Cef; 100 µg/mL), carbendazim (Carb; 40 µg/mL), a combination of Amp, Cef, and Carb at the same concentrations, or no antibiotics. Agar plates were incubated at 25°C under 50 µmol photons m−2 s−1 for 30 days. In parallel, nas-trichome suspensions were cultured in liquid BG11_00_ medium supplemented with Amp, streptomycin (Sm), tetracycline (Tc), chloramphenicol (Cm), nalidixic acid (Nx), or penicillin (Pe) at final concentrations of 10, 30, or 100 µg/mL, and cultures were incubated under the same conditions. Growth of *N. commune* cells and bacteria was assessed in each well after 7 days.

Purification was also attempted by cell sorting using a Cell Sorter SH800S (Sony, Tokyo, Japan). A nas-trichome suspension was streaked onto BG11_00_ agar medium and incubated under the conditions described above. After colonies reached approximately 1 mm in diameter, nas-trichomes were prepared from the colonies and streaked again onto fresh BG11_00_ agar medium. This procedure was repeated three times. A nas-trichome suspension prepared from colonies obtained after the third streak culture was used for cell sorting. Sorting parameters are shown in Table S2. After sorting, the culture fluid was transferred to 384-well plates containing BG11_00_ medium and cultivated as described above. Bacterial growth in each well was monitored during cultivation. *N. commune* colonies for which no bacterial growth was observed were recovered from the wells. Nas-trichome suspensions prepared from these purified colonies were then streaked onto BG11_00_ agar medium and incubated.

**Table S2.**
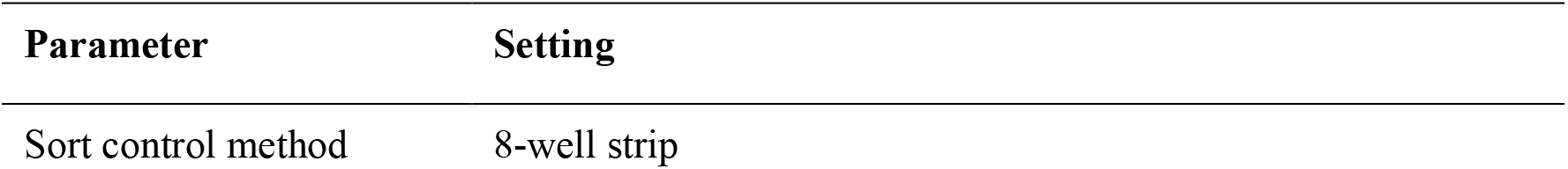
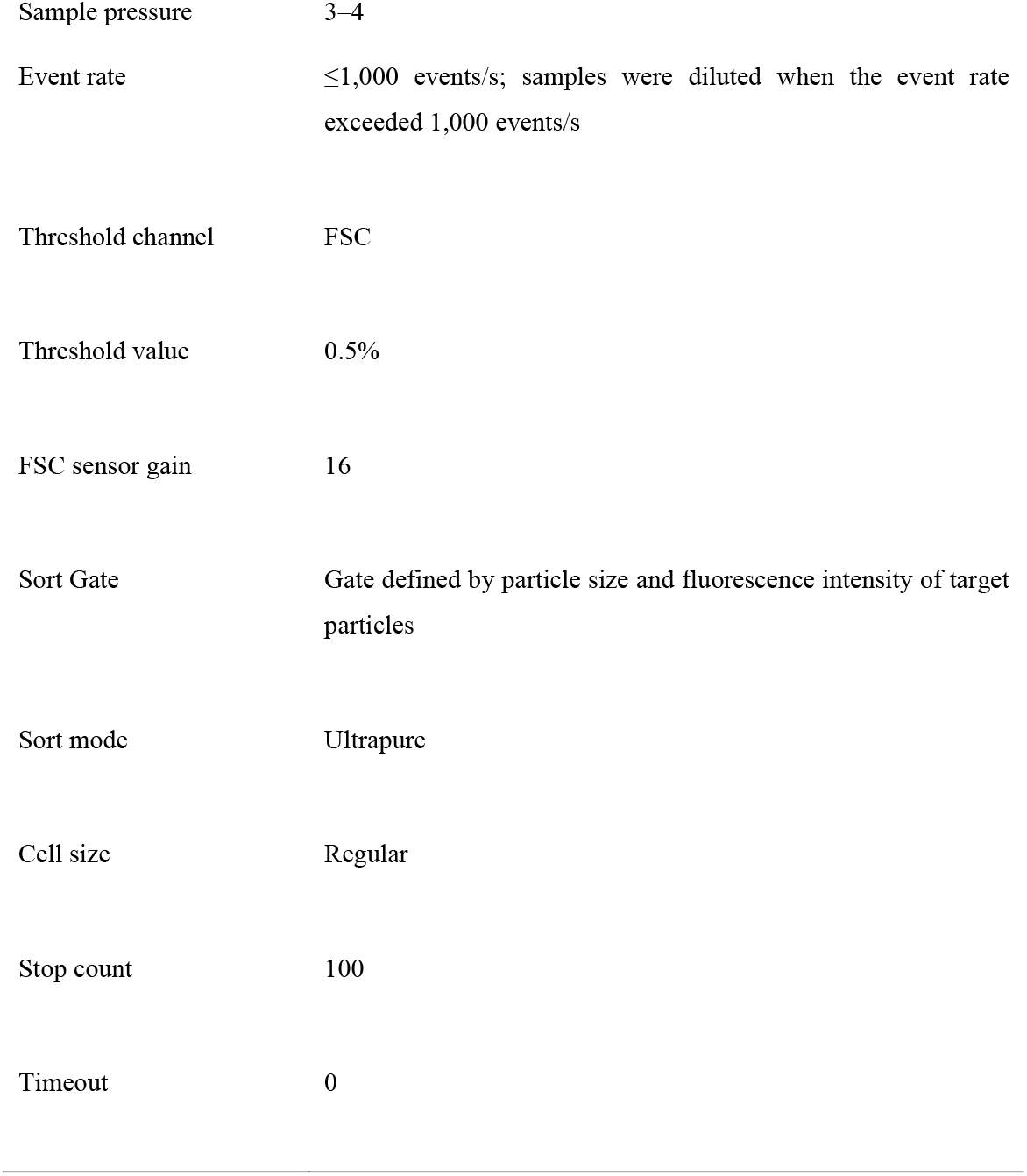
Cell sorter settings used for purification of *Nostoc commune* nas-trichomes.

### Isolation of *N. commune* polysaccharide-assimilating Bacteria

Mibu River colonies were homogenized as described above and dispensed into 96-well plates. Culture fluid was collected from wells in which all *N. commune* cells had died. The suspension was streaked onto *N. commune* polysaccharide agar medium and incubated at 30°C. Resulting bacterial colonies were suspended in DIW, re-streaked onto freshly prepared *N. commune* polysaccharide agar medium, and incubated again at 30°C. Each bacterial suspension was also inoculated into liquid *N. commune* polysaccharide medium and cultured at 30°C. Glycerol stocks were prepared from suspensions showing bacterial growth.

### Co-culture of Purified *N. commune* and Sheath-Formation-Inhibiting Bacteria

One purified *N. commune* colony approximately 5 mm in diameter was transferred to a well of a 48-well plate containing DIW and incubated overnight at 25°C under 50 µmol photons m^−2^ s^−1^. After 2–7 days of incubation, hormogonia released from the colony were collected along with the culture fluid and passed through a 40-µm pluriStrainer to remove colony residues and sheath fragments. The density of the hormogonium suspension was measured using a C-Chip hemocytometer, and the suspension was diluted with DIW to 10 hormogonia per 10 µL. Ten microliters of each sheath-formation-inhibiting bacterial suspension was loaded onto a Bacteria Counting Chamber (depth, 0.02 mm; Erma, Tokyo, Japan), and bacterial cells were counted under a biological microscope. The bacterial suspension was then diluted with DIW to 10^2^, 10^3^, 10^4^, or 10^5^ cells per 10 µL. Each well of a 384-well plate received 50 µL of 10-fold-diluted BG11_00_ liquid medium, 10 µL of the density-adjusted hormogonium suspension, and 10 µL of the bacterial suspension (n = 14 wells per condition). Plates were incubated at 25°C under 50 µmol photons m^−2^ s^−1^. Sheath formation was evaluated on day 7 to assess the effects of candidate sheath-formation-inhibiting bacteria on hormogonium sheath formation.

## Results

### Sheath Formation of *N. commune* Trichomes in 96-Well Plates

As a substitute for hormogonia, which are motile filamentous trichomes released naturally from colonies, nas-trichomes, defined here as non-motile trichome fragments artificially severed from natural colonies, were inoculated into 96-well plates and cultured. During cultivation, the growth morphology of nas-trichomes was classified into two distinct patterns: curling (Fig. 1A, B) and elongation (Fig. 1C, D). Curling occurred when dividing cells remained confined within a sheath and folded into a compact tangle (Fig. 1E). In contrast, when no distinct sheath was observed, trichomes elongated as cell division proceeded (Fig. 1F). Based on these morphological criteria, sheath formation was scored as present or absent. In 96-well plates inoculated with nas-trichomes prepared from colonies subjected to drying and rehydration treatment, wells were clearly classified into curling-rich wells, in which more than 90% of nas-trichomes formed sheaths, and elongation-rich wells, in which more than 90% of nas-trichomes failed to form distinct sheaths. Curling-rich wells were observed in 21 of 36 wells. In contrast, no curling-rich wells were observed when nas-trichomes were prepared from colonies that had not been dried.

Marked bacterial proliferation was frequently observed around nas-trichomes in elongation-rich wells. To examine the relationship between bacterial growth and sheath formation, nas-trichomes prepared from drying- and rehydration-treated colonies were inoculated into 96-well plates again and monitored microscopically for 7 days. Seventy-two nas-trichomes were randomly selected from 12 wells, and the area surrounding each trichome was examined. By day 3, bacteria had proliferated and completely surrounded 55 trichomes, whereas 17 trichomes still retained bacteria-poor areas around them. Among the 55 bacteria-rich trichomes, sheath formation was observed in only 9 trichomes. In contrast, among the 17 bacteria-poor trichomes, 12 formed sheaths (Fig. 2). Furthermore, 31 of the 55 bacteria-rich trichomes were closely associated with dense bacterial aggregates, and only 1 of these 31 trichomes formed a sheath. These results indicate that early bacterial proliferation around nas-trichomes is negatively associated with subsequent sheath formation.

**Fig. 2.**
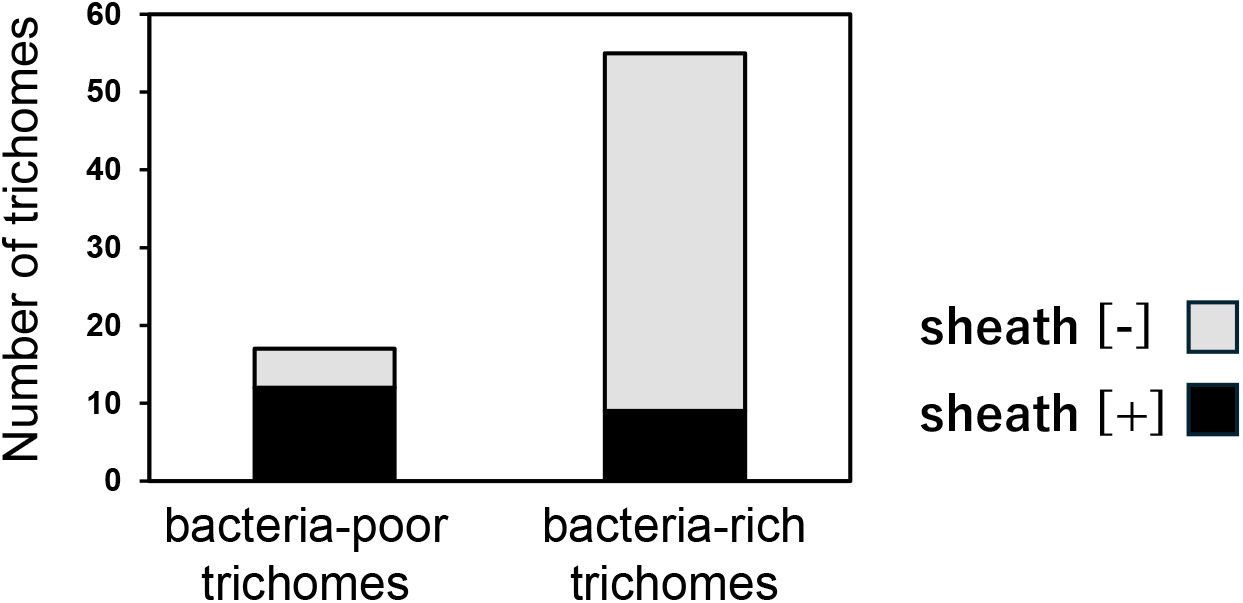
Relationship between bacterial proliferation by day 3 and sheath formation in *N. commune* trichomes. Among 17 bacteria-poor trichomes in which marked bacterial proliferation was not observed by day 3, 12 trichomes formed sheaths (sheath[+], black), whereas 5 did not (sheath[−], gray). Among 55 bacteria-rich trichomes in which bacteria completely surrounded the trichomes by day 3, 9 trichomes were sheath[+] and 46 were sheath[−].

### Bacterial Community Differences between Sheath-Forming and Sheath-Deficient Cultures

Nas-trichome suspensions were prepared again from colonies subjected to drying and rehydration treatment and inoculated into 384-well plates. Based on the proportion of curling-rich and elongation-rich wells, six S(+) groups [S(+)01–06] and eight S(−) groups [S(−)01–08] were selected for 16S rRNA amplicon sequencing. Sequencing of the 14 samples yielded 52,079– 100,021 analyzable reads per sample. Sequences were clustered into operational taxonomic units (OTUs) at 97% similarity, and the number of OTUs ranged from 166 to 652 per sample. OTUs assigned to *Nostoc* were abundant in S(+) samples, accounting for 26–84% of the reads (Fig. 3). In contrast, *Nostoc* OTUs accounted for only 0.7–8.2% of the reads in S(−) samples, whereas OTUs affiliated with *Mucilaginibacter, Ideonella, Blastocatella*, and *Spirosoma* were predominant (Fig. 3). Because the abundance of *N. commune* trichomes was expected to be broadly comparable among samples, the *Nostoc* read count was used as an internal reference to normalize the read counts of non-*Nostoc* taxa. For each sample, a correction factor was calculated by setting the *Nostoc* read count to 100, and this factor was applied to the read counts of all non-*Nostoc* taxa in the same sample. The resulting normalized values were compared between the S(+) and S(−) groups. *Mucilaginibacter, Ideonella*, and *Spirosoma* showed statistically significant differences between the two groups (Fig. 4), identifying these genera as candidates associated with sheath-deficient culture outcomes.

**Fig. 3.**
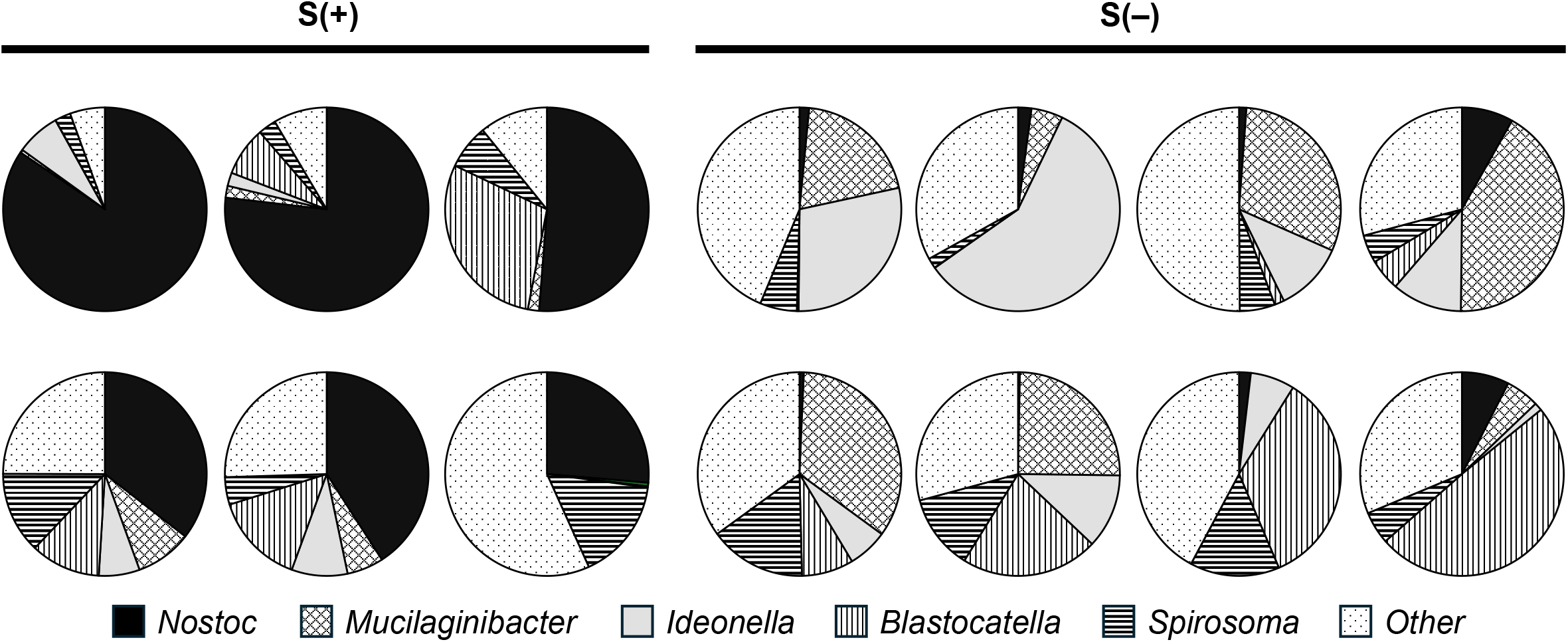
Microbial community composition of culture fluids from S(+) and S(−) groups. For each sample, the proportions of reads assigned to *Nostoc, Mucilaginibacter, Ideonella, Blastocatella, Spirosoma*, and other genera are shown as pie charts. Raw read-count data are provided in Fig. S2.

**Fig. 4.**
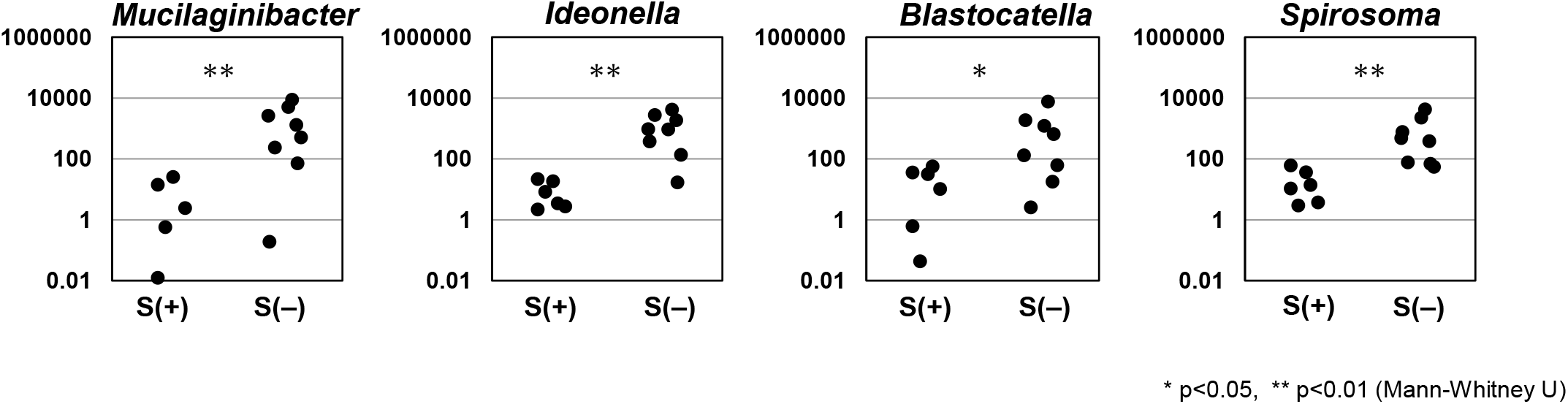
Comparison of normalized abundances of bacterial genera between S(+) and S(−) groups. Normalized abundances of *Mucilaginibacter, Ideonella, Blastocatella*, and *Spirosoma* were calculated by setting the *Nostoc* read count in each sample to 100 and expressing the read counts of each bacterial genus relative to that value. Asterisks indicate statistically significant differences between the S(+) and S(−) groups, as determined by the Mann–Whitney U test: *p ≤ 0.05; **p ≤ 0.01.

To isolate bacteria associated with elongation-rich cultures, S(−) cultures were plated onto *N. commune* polysaccharide agar medium. Four bacterial colonies with distinct colors and morphologies were selected, cultured in liquid *N. commune* polysaccharide medium, and stored as glycerol stocks. These isolates are hereafter referred to as NPA strains. 16S rRNA analysis identified the NPA strains as members of the genera *Methylobacterium, Mucilaginibacter, Spirosoma*, and *Pseudomonas*.

### Purification of *N. commune*

To obtain bacteria-free *N. commune* cultures, we first attempted purification of nas-trichomes by streak culture and antibiotic treatments, as described in Materials and Methods. However, these approaches did not eliminate bacterial contamination while maintaining reproducible growth of *N. commune*. We therefore attempted purification using a cell sorter. Nas-trichomes were separated into microdroplets and analyzed based on light scattering and fluorescence near 660 nm, corresponding mainly to chlorophyll and allophycocyanin. Three particle populations were detected: small particles with weak fluorescence, large particles with strong fluorescence, and particles with intermediate fluorescence and size. Each population was recovered separately and observed microscopically. The population with larger particle size and stronger fluorescence was confirmed to contain *N. commune* nas-trichomes and was subsequently cultured in 384-well plates. Sorted nas-trichomes grew successfully without visible bacterial contamination. Among the sorted nas-trichomes, 65 of 244 formed sheaths, corresponding to a sheath-formation rate of 26.6%, and some developed into spherical colonies more than 5 mm in diameter.

### Density-Dependent Inhibition of Sheath Formation by *Mucilaginibacter* Species

The results described above suggested that bacteria enriched in S(−) samples may inhibit sheath formation in *N. commune* trichomes. To test this possibility, bacteria isolated from S(−) cultures and representative bacterial strains belonging to genera detected in S(−) samples were co-cultured with hormogonia released from purified *N. commune* colonies. The tested strains included the four NPA isolates and reference strains of *Mucilaginibacter* and *Ideonella*. Bacterial suspensions were added at densities of 10^3^–10^5^ cells per well, and sheath formation was evaluated after 7 days. As shown in Fig. 5, *M. segetis, M. oryzae, M. jinjuensis*, and *M. xinganensis* markedly reduced the sheath-formation rate when added at 10^5^ cells per well. In contrast, no significant reduction was observed when these bacteria were added at densities of 10^4^ cells per well or lower. These results demonstrate that specific *Mucilaginibacter* species can inhibit sheath formation in a density-dependent manner, whereas inhibition was not a universal property of all bacteria tested.

**Fig. 5.**
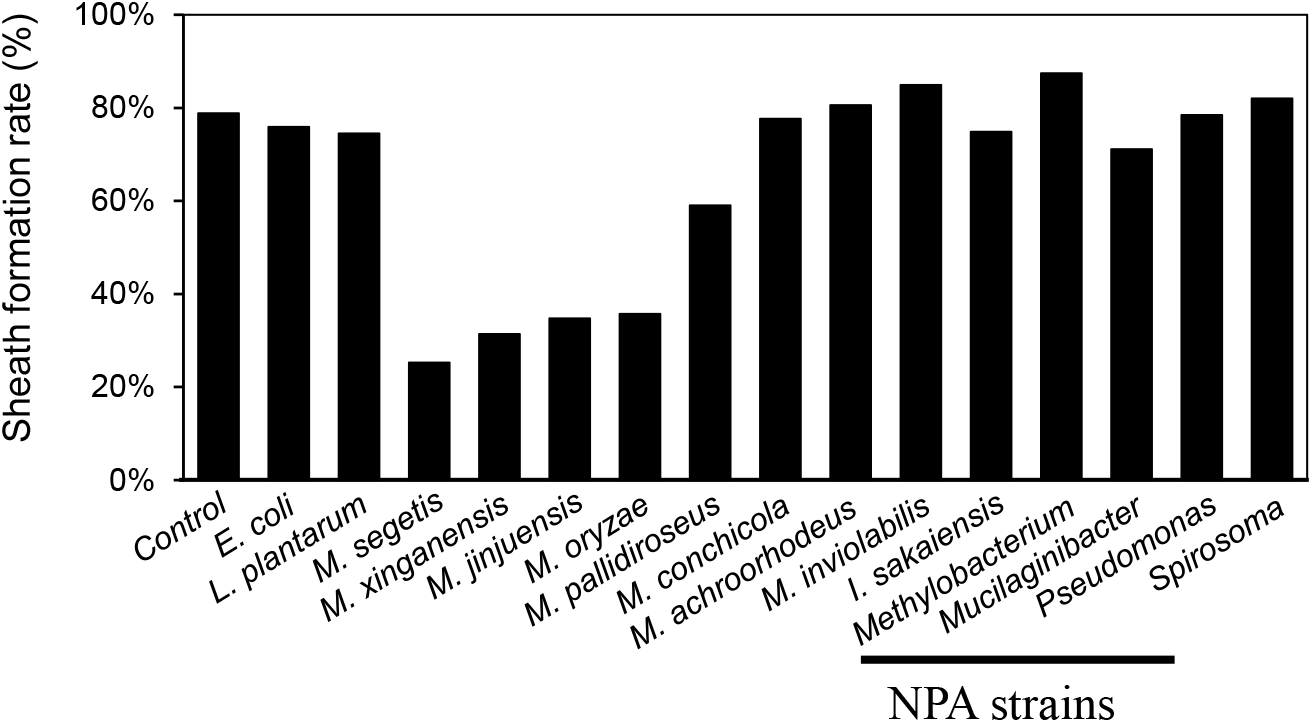
Effects of candidate sheath-formation-inhibiting bacteria on sheath formation in *N. commune* hormogonia. Approximately 10 hormogonia, 10^3^–10^5^ bacterial cells, and culture medium were added to each well of a 384-well plate, with 14 wells per condition. After 7 days of incubation, curling-type trichomes were counted and divided by the total number of trichomes to calculate the sheath-formation rate. Addition of *M. oryzae* (NBRC 106410), *M. jinjuensis* (NBRC 108856), *M. xinganensis* (NBRC 1100384), or *M. segetis* (JCM 34284) significantly reduced the sheath-formation rate at 10^5^ cells per well. No significant reduction was observed at bacterial densities of 10^4^ cells per well or lower.

## Discussion

A reproducible cultivation method for *N. commune* has not yet been established, largely because the environmental and physiological cues that induce hormogonium release remain unclear and because mechanically prepared trichome fragments do not always develop into sheath-enclosed colonies. Under natural conditions, colony propagation appears to involve rupture of the sheath, fragmentation of trichomes near heterocysts, and release of short motile hormogonia, which subsequently initiate new colonies. Because this process cannot currently be induced artificially, mechanical disruption of colonies has been used to generate short, non-motile trichome fragments, referred to here as nas-trichomes. These fragments occasionally develop into sheathed colonies during culture (Katoh et al. 2012). In the present study, microscopic observation revealed two distinct developmental outcomes: a curling type and an elongation type. In the curling type, trichomes became enclosed within a newly formed sheath and folded compactly as cell division proceeded, eventually giving rise to spherical colonies. In contrast, elongation-type trichomes lacked a distinct sheath and extended in a largely filamentous manner while secreting diffuse extracellular material. These morphologies correspond broadly to the developmental forms previously described by Bazzichelli and colleagues, in which microcolonies possess a defined gelatinous sheath, whereas balls of filaments and helical filaments are associated with less distinct extracellular material (Giorgio Bazzichelli 1985, Giorgio Bazzichelli 1986).

A notable feature of these cultures was that individual wells tended to become either curling-rich or elongation-rich, with few intermediate wells containing comparable proportions of both morphologies. This all-or-none pattern suggests that sheath formation was influenced by a factor acting at the well level rather than solely by intrinsic differences among individual trichomes. One plausible explanation is that associated bacteria modulated the early stages of sheath formation. In wells where sheath-degrading or sheath-formation-inhibiting bacteria became dominant, immature extracellular material may have been degraded before a stable sheath could mature, leading to the elongation phenotype. Conversely, when such bacteria remained at low abundance, nas-trichomes may have formed sufficiently robust sheaths to permit curling and compact colony development.

The comparison between drying-treated and non-drying preparations supports this interpretation. Curling-rich wells were observed only when nas-trichomes were prepared from colonies subjected to drying and rehydration, whereas non-dried colonies yielded no curling-rich wells. Drying may have transiently reduced the abundance or activity of bacteria associated with the colony surface or extracellular matrix, thereby allowing sheath formation to proceed during the critical early phase of culture. Consistent with this idea, trichomes that were surrounded by abundant bacteria by day 3 rarely formed sheaths, whereas sheath formation was more frequent when bacterial proliferation remained limited. These observations suggest that the balance between nascent sheath production by *N. commune* and bacterial degradation or interference during the first several days of culture is a key determinant of subsequent morphology.

Microbial community analysis further indicated that members of the genus *Mucilaginibacter* were enriched in elongation-rich cultures. More importantly, co-culture experiments demonstrated that several *Mucilaginibacter* species, including *M. segetis, M. oryzae, M. jinjuensis*, and *M. xinganensis*, significantly reduced sheath formation when introduced at 10^5^ cells per well. This density dependence is consistent with the threshold-like behavior observed in well cultures and supports the view that bacterial abundance during early development determines whether nas-trichomes proceed toward sheath-enclosed curling or sheath-deficient elongation. Notably, most tested bacteria, including several isolates obtained from elongation-rich wells, did not inhibit sheath formation. Thus, the inhibitory activity appears to be restricted to particular bacterial taxa or strains rather than being a general consequence of bacterial coexistence.

An alternative explanation is that bacteria do not directly degrade the immature sheath, but instead alter the physiological state of nas-trichomes through diffusible signals or competition for extracellular metabolites, thereby reducing the ability of *N. commune* to synthesize or organize sheath components. Such an effect could lead to increased leakage of extracellular substances, secondarily promoting bacterial growth in elongation-rich wells. Although this possibility cannot be excluded, the experimental addition of defined *Mucilaginibacter* strains reproduced inhibition of sheath formation, indicating that at least some bacteria can act causally rather than merely proliferating as a consequence of sheath failure.

The mechanism by which these *Mucilaginibacter* species inhibit sheath formation remains to be determined. The genus *Mucilaginibacter* includes species with diverse carbohydrate- or polymer-degrading activities. For example, *M. oryzae* and *M. jinjuensis* have been reported to possess polymer-degrading activities such as cutin-, xylan-, or laminarin-degrading activity (Jeon et al. 2009, Khan et al. 2013), *M. segetis* has been associated with CM-cellulose- and Tween 20-degrading activity (Li et al. 2021), and *M. xinganensis* was described as a phenanthrene-degrading bacterium (Jiang et al. 2019). However, degradation of *N. commune*-derived sheath polysaccharides by *Mucilaginibacter* has not yet been reported. The extracellular polysaccharide of *N. commune* consists of a repeating tetrasaccharide backbone, [→4)-β-D-Glcp-(1→4)-β-D-Xylp-(1→4)-β-D-Glcp-(1→4)-α-D-Galp-(1→], branched at position 6 of a glucose residue by β-linked pyruvated glucuronic acid (Drouillard et al. 2022). To date, an endo-acting enzyme capable of cleaving this polysaccharide into oligosaccharides has been described only from *Runella slithyformis* DSM 19594 and assigned to glycoside hydrolase family GH160 (Helbert et al. 2019). Therefore, identifying enzymes or gene clusters in inhibitory *Mucilaginibacter* strains that are related to GH160-like activities may help clarify the biochemical basis of sheath inhibition.

These findings have practical implications for the cultivation of *N. commune*. Although natural colonies are exposed to diverse microbial communities, sheath formation may proceed successfully when the abundance of inhibitory bacteria remains below a critical threshold. Conversely, culture failure may occur when sheath-formation-inhibiting bacteria proliferate rapidly during the early phase after trichome preparation. Strategies that reduce the initial bacterial load or suppress inhibitory taxa may therefore improve cultivation efficiency. Such strategies could include controlled desiccation pretreatment, optimized washing procedures, selective suppression of sheath-degrading bacteria, or co-culture with non-inhibitory bacteria that competitively limit inhibitory strains without impairing *N. commune* growth. Future studies should identify the enzymatic activities responsible for sheath inhibition, determine whether these activities directly degrade immature sheath polysaccharides, and clarify how microbial community structure influences colony establishment. Overall, this study demonstrates that associated bacteria, particularly several *Mucilaginibacter* species, can inhibit sheath formation in *N. commune* and highlights microbial control as a key requirement for reliable artificial cultivation.

